# Genome analysis of the fatal tapeworm *Sparganum proliferum* unravels the cryptic lifecycle and mechanisms underlying the aberrant larval proliferation

**DOI:** 10.1101/2020.05.19.105387

**Authors:** Taisei Kikuchi, Mehmet Dayi, Vicky L. Hunt, Atsushi Toyoda, Yasunobu Maeda, Yoko Kondo, Belkisyole Alarcon de Noya, Oscar Noya, Somei Kojima, Toshiaki Kuramochi, Haruhiko Maruyama

**Affiliations:** Faculty of Medicine, University of Miyazaki, Miyazaki, 889-1692 Japan; Forestry Vocational School, Duzce University, 81620, Duzce, Turkey; Department of Biology and Biochemistry, University of Bath, Bath, BA27AY UK; Comparative Genomics Laboratory, Department of Genomics and Evolutionary Biology, National Institute of Genetics, Mishima, Shizuoka, 411-8540 Japan; Division of Medical Zoology, Department of Microbiology and Immunology, Faculty of Medicine, Tottori University, Yonago 683-8503, Japan; Institute of Tropical Medicine, Central University of Venezuela, Venezuela, 2101 Maracay, Caracas, Venezuela; Department of Clinical Laboratory Medicine, Chiba-Nishi General Hospital, Matsudo City, Chiba, 270-2251 Japan; Department of Zoology, National Museum of Nature and Science, 4-1-1 Amakubo, Tsukuba, Ibaraki, 305-0005 Japan

**Keywords:** pseudophyllidean tapeworm, gene family evolution, relaxed selection, extracellular matrix coordination, asexual reproduction, oncogenes, homeobox, fibronectin, cadherin

## Abstract

**Background:** The cryptic parasite *Sparganum proliferum* proliferates in humans and invades tissues and organs. Only scattered cases have been reported, but *S. proliferum* infection is always fatal. However, the *S. proliferum* phylogeny and lifecycle are still an enigma.

**Results:** To investigate the phylogenetic relationships between *S. proliferum* and other cestode species, and to examine the underlying mechanisms of pathogenicity, we sequenced the entire *S. proliferum* genome. Additionally, *S. proliferum* plerocercoid larvae transcriptome analyses were performed to identify genes involved in asexual reproduction in the host. The genome sequences confirmed that the *S. proliferum* genetic sequence is distinct from that of the closely related *Spirometra erinaceieuropaei*. Moreover, nonordinal extracellular matrix coordination allows for asexual reproduction in the host and loss of sexual maturity in *S. proliferum* is related to its fatal pathogenicity in humans.

**Conclusions:** The high-quality reference genome sequences generated should prove valuable for future studies of pseudophyllidean tapeworm biology and parasitism.

## Background

The cryptic parasite *Sparganum proliferum* was first identified in a 33-year-old woman in Tokyo in 1904. The patient’s skin was infected with numerous residing cestode larva of unknown taxonomy. Ijima *et al* [1] originally designated the parasite as *Plerocercoides prolifer*, and considered it a pseudophyllidean tapeworm in the plerocercoid larval stage. In 1907, an extremely similar human case was reported by Stiles in Florida, USA, and the responsible parasite was renamed *S. proliferum* [2]. Clinical symptoms and post-mortem findings indicate that *S. proliferum* proliferates in humans and invades various organs and tissues, including the skin, body walls, lungs, abdominal viscera, lymph nodes, blood vessels, and the central nervous system, leading to miserable disease prognosis [3, 4]. Not many cases have been reported to date but the infection was fatal in all reported cases (reviewed in [5]).

There was a postulation about the origin of this parasite. Some parasitologists considered it to be a new species of pseudophyllidean tapeworm, whereas others suspected that *S. proliferum* was a virus-infected or aberrant form of *Spirometra erinaceieuropaei*, based on morphological similarities [6, 7]. Recent DNA sequence analyses of mitochondrial NADH dehydrogenase subunit III, mitochondrial tRNA, cytochrome oxidase subunit I, and nuclear succinate dehydrogenase iron-sulfur protein subunit (sdhB) genes suggested that *S. proliferum* is a closely related but distinct species of *S. erinaceieuropaei* [8, 9]. However, the adult stage of *S. proliferum* has not been observed and the precise taxonomic relationships of *S. proliferum* with other worms remain unclear because few genes have been sequenced.

In addition to taxonomic considerations, the pathogenicity of *S. proliferum* and its mechanisms of proliferation and invasion in mammalian hosts are of considerable interest. In principle, plerocercoids of pseudophyllidean tapeworms (spargana), including those of *S. erinaceieuropaei* and other *Spirometra* species, do not proliferate asexually, but migrate through subcutaneous connective tissues, causing only non-life threatening sparganosis (non-proliferative sparganosis). Other organs, such as the lungs and liver or the central nervous system, may be niches for these worms but are not commonly described. Symptoms of non-proliferative sparganosis are mainly caused by the simple mass effect [5].

Asexual proliferation of larvae and the destruction of host tissues are characteristic of cyclophyllidean tapeworms such as *Echinococcus*, which proliferates asexually by generating a peculiar germinative layer in a hydatid cyst form [10]. In another cyclophyllidean tapeworm, *Mesocestoides*, asexual multiplication is achieved by longitudinal fission [10, 11]. In contrast, the pseudophyllidean *Sparganum* plerocercoid undergoes continuous branching and budding after invading the human body by an unidentified route, and produces vast numbers of progeny plerocercoids.

To clarify the phylogenetic relationship of the enigmatic parasite *S. proliferum* with other cestode species and investigate the underlying pathogenic mechanisms, we sequenced its entire genome as well as that of newly isolated *S. erinaceieuropaei*. We also performed transcriptome analyses of *S. proliferum* plerocercoid larvae to identify genes that are involved in asexual reproduction in the host. Those analyses revealed its phylogeny and gene evolution that contribute to the proliferation and pathogenicity of *S. proliferum*.

## Results

### Genomic features of *S. proliferum* and *S. erinaceieuropaei*

We sequenced the *S. proliferum* genome using multiple insert-length sequence libraries (Additional Table S1) and compiled a 653.4-Mb assembly of 7388 scaffolds with N50 of 1.2 Mb. The *S. erinaceieuropaei* genome was assembled into 796 Mb comprising 5723 scaffolds with N50 of 821 kb. These assembly sizes were 51.9% and 63.2% of the previously published *S. erinaceieuropaei* genome (UK isolate) [12]. CEGMA and BUSCO report the percentage of highly conserved eukaryotic gene families that are present as full or partial genes in assemblies and nearly 100% of core gene families are expected in most eukaryote genomes. BUSCO analyses showed that 88.1% and 88.5% of core gene families were represent in *S. proliferum* and *S. erinaceieuropaei* genomes, respectively, higher than or comparable to other previously published tapeworm genomes (Table 1). CEGMA completeness values for *S. proliferum* and *S. erinaceieuropaei* were slightly lower than those from BUSCO analyses. Low CEGMA completeness was also seen in other pseudophyllidea tapeworm genomes, including *S. erinaceieuropaei* UK isolate, *Diphyllobothrium latum*, and *Schistocephalus solidus* (Table 1). Low CEGMA completeness values of these two genome assemblies, therefore, indicate pseudophyllidean-specific loss or high divergence of the genes that are conserved in other eukaryotic taxa. The average numbers of CEGs (hits for 248 single-copy eukaryotic core genes) for *S. proliferum* and *S. erinaceieuropaei* were 1.2 and 1.3, respectively, indicating that the assembly sizes roughly represent the haploid genome sizes of these tapeworms. However, in K-mer analyses of Illumina short reads, we estimated haploid genome sizes of 582.9 and 530.1 Mb for *S. proliferum* and *S. erinaceieuropaei*, respectively (Additional Fig S1a), indicating that the assemblies contain heterozygous haplotypes and/or overestimated gap sizes. Ploidies were inferred from heterozygous K-mer pairs and were diploid for both species (Figure S1b).

**Table 1.** Statistics of genome assemblies

|  | *Sparganum proliferum* (v2.2) | *Spirometra erinaceieuropaei* (v2.0) | *Spirometra erinaceieuropaei* (UK) (WBPS13) | *Diphyllobotrium latum* (WBPS13) | *Schistocephalus solidus* (WBPS13) | *Hymenolepis microstoma* (WBPS13) | *Taenia solium* (WBPS13) | *Echinococcus multilocularis* (WBPS13) |
| --- | --- | --- | --- | --- | --- | --- | --- | --- |
| Assembly size (Mb) | 653.4 | 796.0 | 1258.7 | 531.4 | 539.4 | 182.1 | 122.4 | 114.5 |
| Num. scaffolds | 7,388 | 5,723 | 482,608 | 140,336 | 56,778 | 3,643 | 11,237 | 1,288 |
| Average (kb) | 88.4 | 139.0 | 2.6 | 3.8 | 10.0 | 50.0 | 11.2 | 889.3 |
| Largest scaff (kb) | 8,099 | 5,490 | 90 | 80 | 595 | 2,234 | 740 | 15,981 |
| N50 (kb) | 1,242 | 821 | 5 | 7 | 32 | 767 | 68 | 5,229 |
| N90 (kb) | 110 | 167 | 1 | 2 | 5 | 41 | 5 | 213 |
| Gaps (kb) | 51,020 | 77,788 | 128,163 | 38,407 | 22,091 | 259 | 164 | 336 |
| CEGMA completeness complete/partial (%) | 61.7/81.5 | 58.5/80.2 | 29.4/45.9 | 49.6/65.3 | 76.6/87.9 | 91.9/92.7 | 87.1/90.7 | 93.2/93.2 |
| Average CEG number complete/partial | 1.1/1.2 | 1.1/1.3 | 1.8/2.2 | 1.5/1.6 | 1.2/1.3 | 1.1/1.1 | 1.2/1.2 | 1.1/1.1 |
| BUSCO completeness (Metazoa dataset/Eukaryota dataset) | 72.0/88.1 | 71.9/88.5 | 33.6/37.3 | 38.1/53.8 | 70.3/86.2 | 78.6/90.4 | 72.6/85.5 | 72.2/88.1 |
| Num. coding genes | 25,627 | 30,751 | 39,557 | 19,966 | 20,228 | 12,373 | 12,481 | 10,273 |
| Coding gene size (median; aa) | 665.0 | 627.0 | 200.0 | 216.0 | 455.0 | 709.0 | 609.0 | 596.0 |

The genomes of *S. proliferum* and *S. erinaceieuropaei* are highly repetitive, with about 55.0% repetition of the total genome length in both genomes (Additional Fig S2 and Table 2). Long interspersed nuclear elements (LINEs) occupy 26.3% and 31.9% of the total genomes of *S. proliferum* and *S. erinaceieuropaei*, respectively. These LINEs predominantly comprise the three types (Penelope, RTE-BovB, and CR1), which are also abundant in other pseudophyllidea genomes (Additional Fig S2).

**Table 2.**
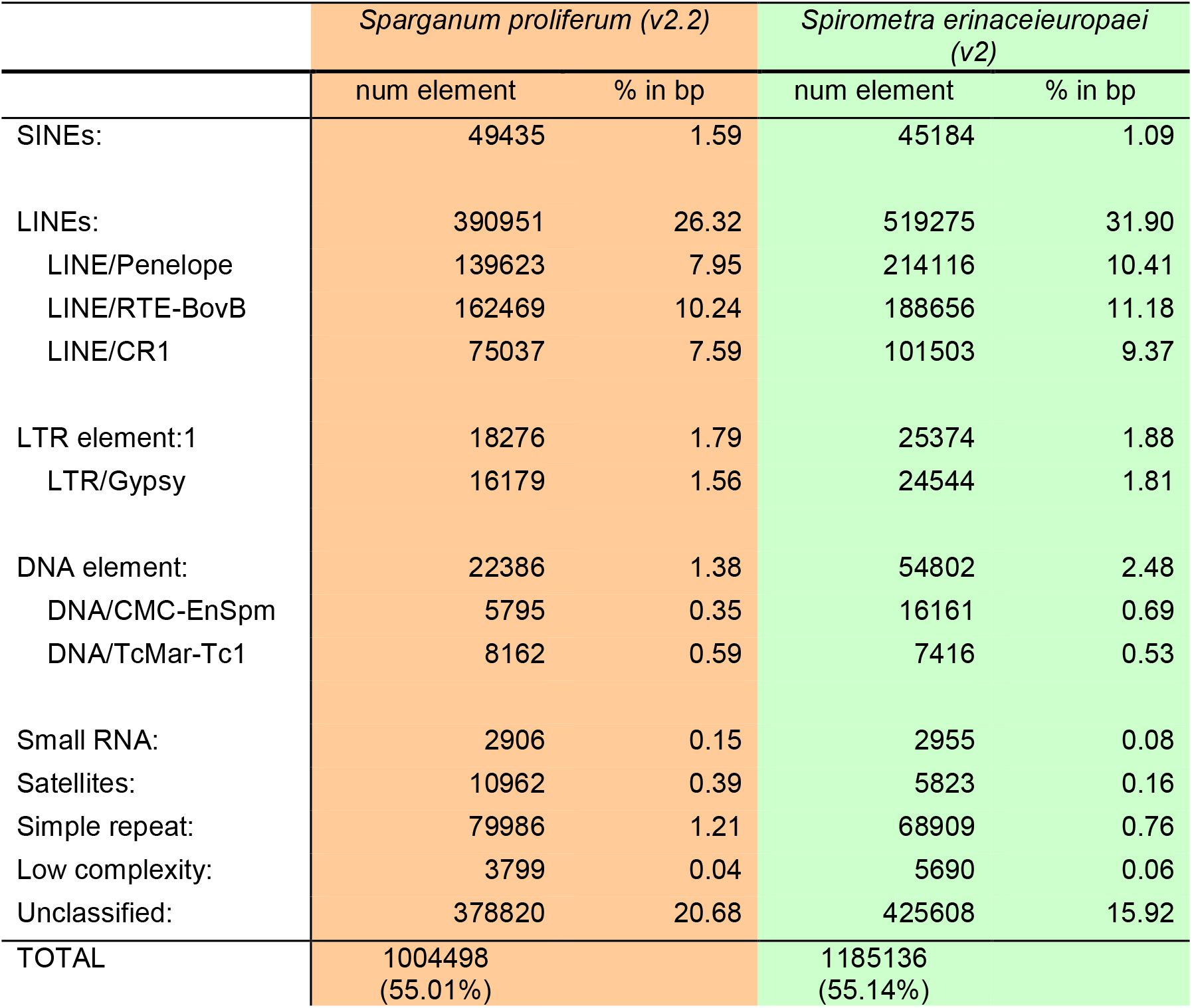
Statistics of repeats in genomes

A total of 25627 genes were predicted in *S. proliferum* assemblies, about 5000 fewer than for *S. erinaceieuropaei* (30751), but more numerous than for other cestode genomes. In studies of the *S. erinaceieuropaei* UK isolate {Bennett, 2014 #39}, the gene number (> 39000) was likely overestimated due to fragmentation and redundancy in the assembly.

### Phylogenetic placement of *S. proliferum*

Phylogenetic relationships of *S. proliferum* with other cestode species were inferred from 205 single-copy orthologues (Figure 1). A clear separation was identified between pseudophyllidea and cyclophilidea clades. In the pseudophyllidea clade, *S. proliferum* occupied the basal position of the *Spirometra* cluster, in which two *S. erinaceieuropaei* isolates (Japan and UK isolates) were placed beside each other.

**Figure 1.**
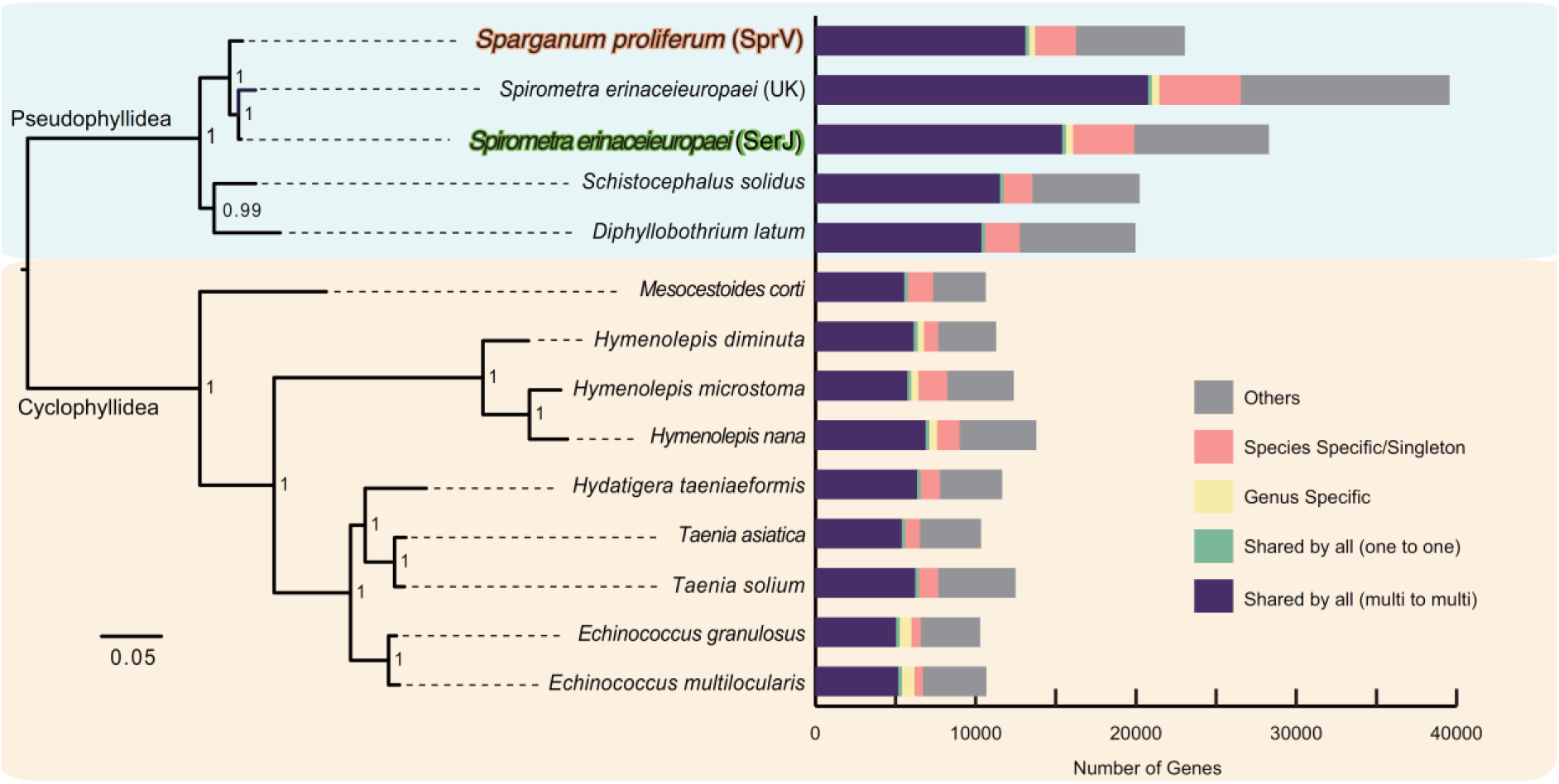
Phylogeny and gene contents; genes are categorized in a stack bar, and the length of stack bar is proportional to number of genes.

Phylogenetic tree topology based on mitogenomes of the 14 cestodes and all available mitogenome data of *Spirometra* in the GenBank, was similar to that of the nuclear genome (Additional Fig S3). Yet in contrast with the nuclear genome tree, the *S. erinaceieuropaei* UK isolate was located in a basal position of the *Spirometra* cluster, placing *S. proliferum* in the middle of *Spirometra* species, albeit with a long branch. These inconsistencies between nuclear and mitogenome trees may reflect uncertainties of species classification in the genus *Spirometra* [13, 14]. Moreover, mitochondrial sequences can give poor inferences of species trees [15]. Cumulatively, these results suggest that *S. proliferum* has a close phylogenetic relationship with *Spirometra* but is clearly distinguished by genomic features and gene contents.

### Gene family evolution

Protein family (Pfam) analyses revealed highly similar protein domain distributions of *S. proliferum* and *Spirometra* genomes (r = 0.99; Figure 2, Additional Table S2). Few domains differed significantly in abundance between the two species. Among these, the *S. proliferum* genome was underrepresented in zinc-finger families (Zf-C2H2, Zf-C2H2_4, Zf-C2H2_6, Zf-C2H2_jaz and Zf-met), reverse transcriptase (RVT_1), exo/endonuclease/phosphatase, galactosyltransferase, and alpha/beta hydrolase (abhydrolase_6). Overrepresented Pfam domains in *S. proliferum* included a distinct type of zinc-finger domain (zf-3CxxC), fibronectin type III (fn3), trypsin, RNA polymerase III RPC4 (RNA_pol_Rpc4), and an ADP-specific phosphofructokinase/glucokinase conserved region (ADP_PFK_GK).

**Figure 2.**
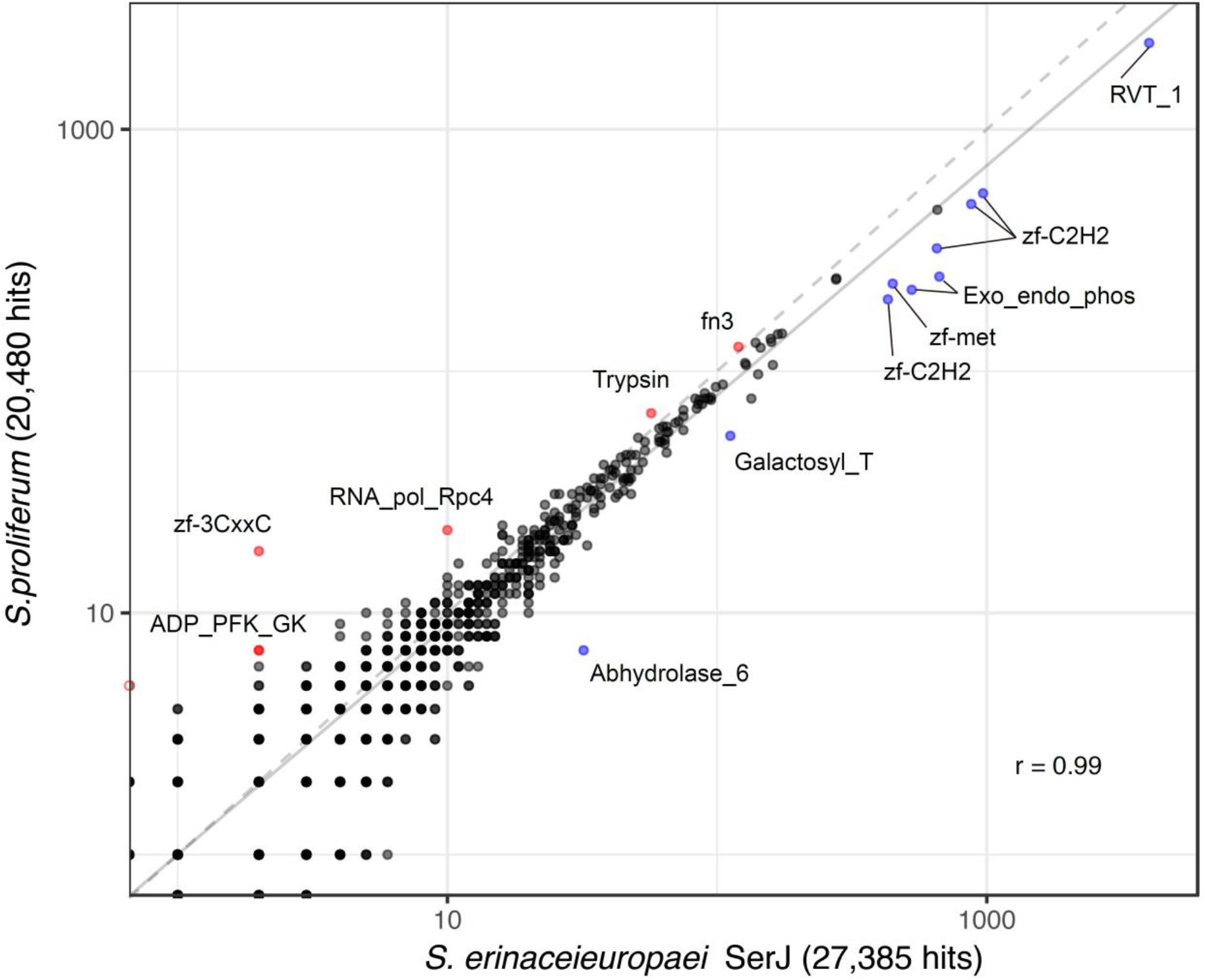
A scatterplot showing the abundance of Pfam domains in *S. proliferum* and *S. erinaceieuropaei* genomes; Pfam domains that are more enriched in *S. proliferum* than in *S. erinaceieuropaei* are highlighted in red. Those enriched in *S. erinaceieuropaei* relative to *S. proliferum* are highlighted in blue.

We performed gene family analysis using OrthoFinder with the predicted proteomes of *S. proliferum*, *S. erinaceieuropaei*, and other selected cestode genomes. A total of 234522 proteins from 14 cestode species were placed into 39174 gene families (Figure 1). The *S. proliferum* proteome (25627 proteins) was encoded by 9136 gene families, among which 7364 were shared by all 14 cestodes and 2550 proteins were specific to the species or singleton. The *S. erinaceieuropaei* proteome (30751 proteins) was clustered into 9008 gene families, 3806 of which were species specific or singletons. Only four gene families were specific to both *Spirometra* and *Sparganum*.

We used computational analysis of gene family evolution (CAFE) to estimate gene family expansion and contraction, and identified gene families with significantly higher than expected rates of gains and losses (Figure 3, Additional Table S3). Twenty-one gene families were significantly expanded in the *S. proliferum* lineage, and these included annotations for fibronectin, reverse transcriptase, zinc-finger C2H2 type, and core histone (Additional Table S4). Significantly contracted gene families (43 families) had annotations relating to signal transduction proteins, such as phosphatases and kinases, and ion channels and ABC transporters (Additional Table S5). Fibronectin, reverse transcriptase, zinc-finger C2H2 type, and peptidases were present in expanded and contracted families.

**Figure 3.**
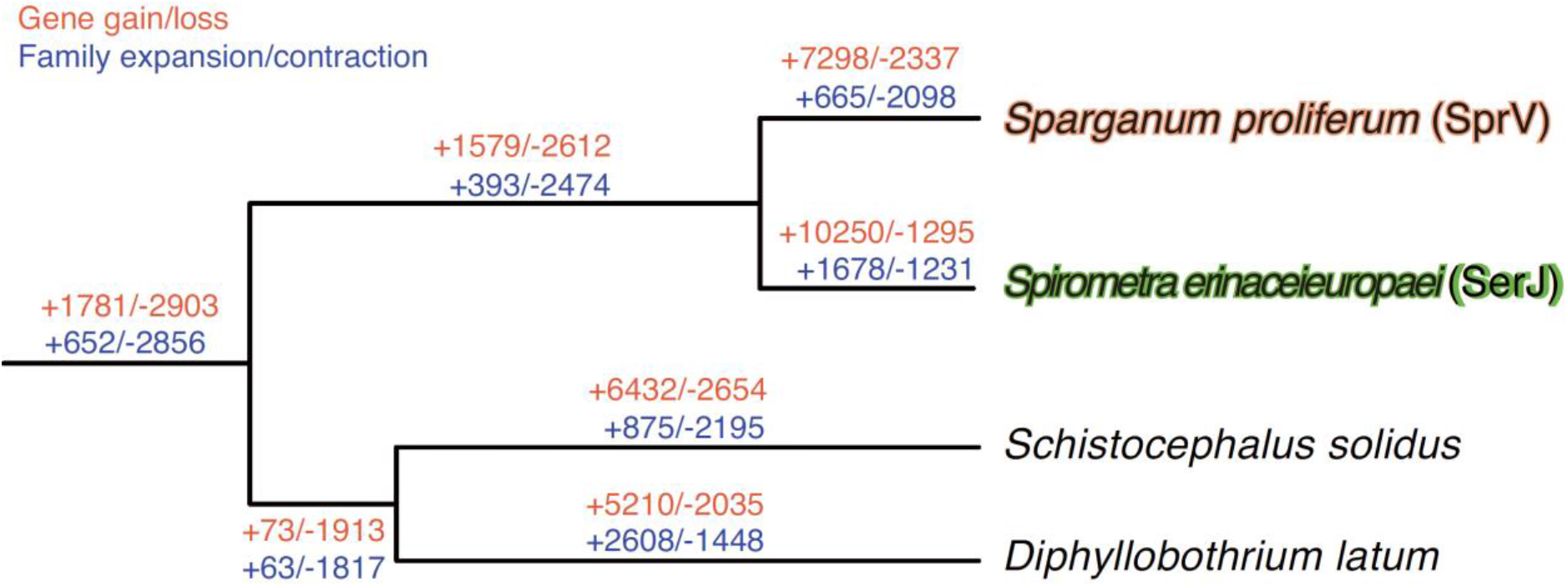
Gene family evolution of selected cestode species was inferred using computational analysis of gene family evolution (CAFE). Numbers on each branch (or lineage) indicate specific gains/losses of that branch (or lineage).

In the *S. erinaceieuropaei* lineage, 63 and 15 gene families were significantly expanded or contracted (Additional Table S6 and S7), respectively. Among them, highly lineage specific expansion was found for 7 families (i.e. 10 or more genes in *S. erinaceieuropaei*, whereas one or no genes in *S. proliferum*. For example, the Orthogroup OG0000184 contains one *S. proliferum* gene and 44 *S. erinaceieuropaei* genes, encoding biphenyl hydrolase-like protein (BPHL), which harbors the Pfam domain abhydrolase_6 (Figure 2). Although the other gene families mostly encode proteins of unknown function, they were likely expanded after speciation from *S. proliferum* and *S. erinaceieuropaei* and may have specific roles in the *S. erinaceieuropaei* lifecycle or parasitism.

### Conserved developmental pathway genes

Homeobox transcription factors are involved in patterning of body plans in animals. The homeobox gene numbers are much fewer in parasitic flatworms than in most other bilaterian invertebrates, which have a conserved set of approximately 100 homeobox genes. Genome severance of four cyclophyllid cestodes revealed that out of 96 homeobox gene families that are thought to have existed at the origin of the bilateria, 24 are not present in cestodes [16]. The pseudophyllid cestodes *S. proliferum* and *S. erinaceieuropaei* have similar homeobox class repertoires as those in cyclophyllid cestodes, in which class ANTP was the most abundant, followed by classes PRD and TALE; Table 3). The total numbers of homeobox domains identified in *S. proliferum* and *S. erinaceieuropaei* are 64 and 71, respectively, and because these were fewer than in the cyclophyllids *Echinococcus multilocularis* and *Taenia solium* (Table 3), they are the most reduced of any studied bilaterian animal. The three homeobox families Pou/Pou6, ANTP/Bsx, and ANTP/Meox were not present in *S. proliferum* and *S. erinaceieuropaei*, whereas the homeobox family ANTP/Ro was found in *S. proliferum* and *S. erinaceieuropaei* but not in *E. multilocularis* and *T. solium* (Additional Fig S4).

**Table 3.** Homeobox complement in *S. proliferum* and *S. erinaceieuropaei* compared with other tapeworms and bilaterians

| homeobox class | *Sparganum*<br>*proliferum*<br>(v2.2) | *Spirometra*<br>*erinaceieuropaei*<br>(v2.0) | *Taenia solium*<br>(WBPS13) | *Echinococcus*<br>*multilocularis*<br>(WBPS13) | *Branchiostoma*<br>*floridae* |
| --- | --- | --- | --- | --- | --- |
| ANTP | 25 | 30 | 36 | 25 | 58 |
| PRD | 10 | 8 | 11 | 15(18) | 21 |
| CUT | 3 | 4 | 3 | 3 | 4 |
| SINE | 3 | 4 | 2 | 3 | 3 |
| TALE | 8 | 10 | 11 | 12 | 10 |
| CERS | 2 | 1 | 2 | 2 | 1 |
| POU | 3 | 4 | 4 | 5 | 6 |
| LIM | 6 | 6 | 7 | 7 | 8 |
| ZF | 4 | 4 | 3(2) | 3(2) | 4 |
| Total | 64 | 71 | 79 | 75 | 115 |

Comparisons between *S. proliferum* and *S. erinaceieuropaei* showed that the homeobox families TALE/Pknox, ANTP/Hox1, ANTP/Msxlx, and POU/Pou-like are missing in *S. proliferum*, despite being present in the other cestodes. In contrast, the homeobox families ANTP/Dbx and PRD/Alx were found in *S. proliferum* but not in *S. erinaceieuropaei*.

Other conserved genes with roles in flatworm developmental pathways, such as Hedgehog and Notch, were conserved in *S. proliferum* and *S. erinaceieuropaei*. But in the Wnt pathway, whose complement is much smaller than the ancestral complement in tapeworms [16], two further genes (Axin and LEF1/TCF) were missing in *S. proliferum* and *S. erinaceieuropaei* (Table S8).

### Horizontally transferred genes

To determine whether the present genomes contained horizontally transferred genes (HTGs) from other organisms, we used a genome-wide prediction method based on a lineage probability index using the software Darkhorse2 identified 19 and 33 putative HTGs in *S. proliferum* and *S. erinaceieuropaei*, respectively (Additional Table S9 and S10). For these transfers, all possible host organisms were bacteria except for one *Spirometra* gene that has high similarity to a chlorella virus gene. Orthologues of most *S. proliferum* putative HTGs were also detected as horizontally transferred in *S. erinaceieuropaei*. Moreover, possible host bacteria, including *Marinifilum breve*, *Aphanizomenon flos-aquae, Alcanivorax* sp., and *Vibrio* sp., were shared by the two cestode species and were aquatic or marine bacteria, indicating that these genes were acquired by a common ancestor of the two tapeworms which had aquatic phase in the life cycle.

### Positive selection of the *S. proliferum* lineage

Positive selection is a mechanism by which new advantageous genetic variants sweep through a population and drive adaptive evolution. To investigate the roles of positive selection in the evolution of *S. proliferum*, we performed dN/dS branch-site model analyses with single-copy orthologous genes from 12 tapeworms and identified a total of 35 positively selected genes in the *S. proliferum* lineage (Additional Table S11). Evolutionary pressures were identified for some genes that are essential to cellular processes, including transcription/RNA processing/translation genes encoding DNA-directed RNA polymerase II subunit, polypyrimidine tract-binding protein, adenylate kinase, ribosomal protein L21, snu13 NHP2-like protein, and eukaryotic translation initiation factors. Other identified genes were related to transportation (dynein intermediate chain 2) and mitochondrial processes (Rieske). Genes involved in stress and immune responses, such as DNAJ/Hsp40, HIKESHI protein, Toll-like receptor, and Ig_3/Ig, were also positively selected in the *S. proliferum* lineage, along with the RAS oncogene *Rab-4A*.

Environmental change often eliminates or weakens selective pressures that were formerly important for the maintenance of a particular trait [17]. We detected 9 genes that were subject to these circumstances of “relaxed selection” in the *S. proliferum* lineage, relative to the other tapeworm lineages (Additional Table 12). These genes encode proteins with putative roles in developmental regulation and cell differentiation. In particular, the receptor roundabout (ROBO) and secreted molecules of the SLIT family, together, play important roles in guiding axons and proper morphogenesis [18]. The Rho GTPase-activating protein is also highly expressed in highly differentiated tissues and affects cell differentiation by negatively regulating Rho-GTPase signaling [19]. Delta-like protein (DLL) is an inhibitory ligand of the Notch receptor pathway and is expressed during brain development [20]. Vascular endothelial growth factor receptor is also known to regulate stem cell homeostasis and repopulation in planarian species [21]. Hence, these instances of relaxed selection indicate that the worm has long since used certain developmental pathways. We also identified two genes encoding cadherin (protocadherin) that were subject to relaxed selection. Cadherein is a transmembrane protein that mediates cell–cell adhesion in animals and those relaxed selections indicate diverging cell adhesion process in the worm.

### Differential gene expression involved in asexual proliferation and parasitism

We maintained *S. proliferum* via serial infection of mice and found that some plerocercoid worms exhibit a highly branching structure (medusa-head form; Figure 4a), which was observed frequently in heavily infected mice. In contrast, in mice with low worm burdens, most worms had unadorned non-branching morphology (wasabi-root form). Worms with the medusa-head form are considered the main sources of new plerocercoid worms in the host, and their proliferation is highly related to their pathogenicity. We, therefore, identified genes with expression levels that distinguished medusa-head and wasabi-root forms.

**Figure 4.**
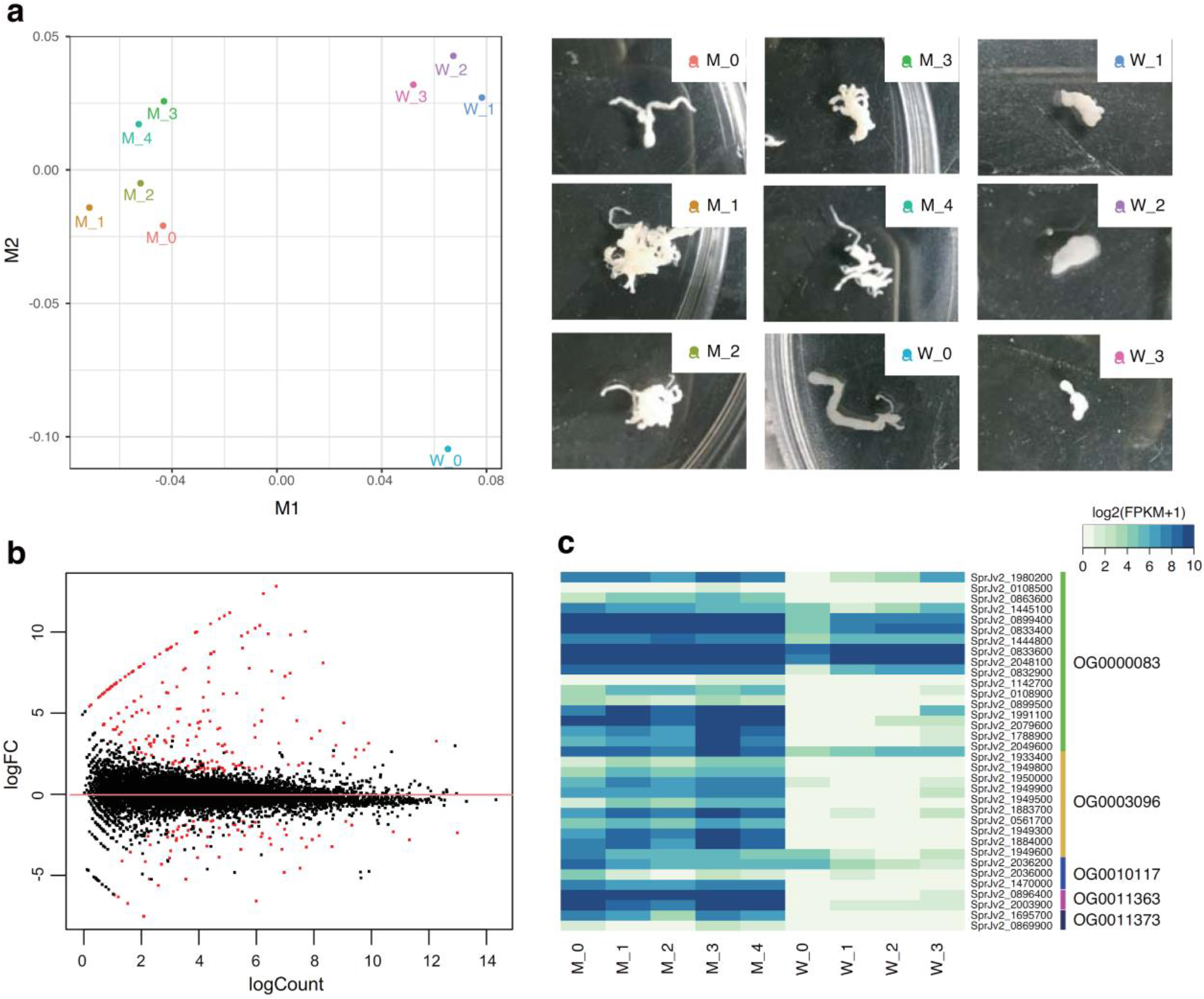
Comparison of gene expression in highly branching worms (medusa-head form) relative to static worms (wasabi-root form) of *S. proliferum*: A) multidimensional scaling (MDS) analyses of RNA-seq samples clearly separate the two forms by dimension 1. Pictures of used samples are shown on the right. B) Bland-Altman (MA) plot of the two-form comparison; dots represent transcripts and log2 fold changes (medusa-head/wasabi-root) plotted against average abundance in counts per million. Red dots indicate differentially expressed transcripts with false discovery rates (FDR) of < 0.05 and fold changes of > 2. C) Heatmap of gene families encoding novel secreted proteins; the heat map shows log2 fragments per kilobase per million reads mapped (FPKM) values for 5 gene families.

RNAseq analysis revealed 357 differentially expressed genes (DE genes) between medusa-head and wasabi-root forms (246 upregulated and 111 downregulated in medusa-head) (Figure 4b). The upregulated set in medusa-head forms were dominated by genes encoding peptidases and peptidase inhibitors, such as tolloid-like proteins (19 genes), chymotrypsin-like proteins (6 genes) and CAP domain-containing proteins (12 genes) as well as transposon-related proteins such as gag-pol polyproteins and reverse transcriptases (30 genes) (Additional Table S13). This set of DE genes was enriched in the GO categories for metalloendopeptidase activity and proteolysis (Additional Table S14). Downregulated genes also encoded a variety of peptidases and peptidase inhibitors, including leucyl aminopeptidase (5 genes), chymotrypsin-like elastase (7 genes), and kunitz bovine pancreatic trypsin inhibitor domain protein (3 genes), with high representation under the GO terms metalloexopeptidase, aminopeptidase, and manganese ion binding (Additional Table S14). Peptidases and peptidase inhibitors are secreted by many types of pathogens, including bacteria, fungi, and parasites, and often play critical roles in survival and virulence [22–24].Other genes known to be involved in pathogenicity in other pathogens were also upregulated in the medusa-head form, including genes encoding multidrug resistance-associated proteins [25] and tetraspanins. The latter proteins have four transmembrane domains and not only play roles in a various aspects of cell biology but also are used by several pathogens for infection and regulate cancer progression [26].

Genes that are involved in cell-growth and cancer development were also upregulated in the medusa-head form, including those encoding proteins from wnt (wnt-111 and wnt-5) and ras/rab (ras-0b, ras-2 and Rasef) pathways, transcription factors/receptors (sox1a, fibroblast growth factor receptor) and homeobox proteins (prospero, PAX, orthopedia ALX and ISL2).

It has been shown that expansions of gene families and changes in expression levels have been associated with the evolution of parasitism in previous studies [27, 28]. An upregulation of genes from expanded gene families was also found in *S. proliferum*. For instance, 15 genes were identified as upregulated from an expanded gene family (OrthoGroup OG0000040). The orthogroup OG0000044 includes genes encoding mastin precursors, and six of these were upregulated and another six were downregulated in the medusa-head form (Additional Table S13). Phylogenetic analyses of those gene families indicate that some of these orthogroups are conserved across flatworms, while others are specific to the Pseudophillidea clade of flatworms (Additional Fig S5).

Among the present DE genes, 85 that were upregulated in medusa-head forms have no known functions. These included 17, 10, 3, 2, and 2 genes from orthogroups OG0000083, OG0003096, OG0010117, OG0011363, and OG0011373, respectively. These orthogroups were expanded in the *S. proliferum* lineage and the DE genes had extremely high fold changes (Figure 4c). Because their products predominantly harbour secretion signal peptides (Additional Table S13), they are likely to be secreted by the parasite into the host and play important roles in parasitism, aberrant larval proliferation in the host, and/or modulation of host immunity.

## Discussion

*S. proliferum* is a cryptic parasite with fatal consequences, but its phylogeny and lifecycle are poorly understood. In this study, we sequenced the *S. proliferum* genome and performed comparative genomics with other tapeworm species, including the newly-sequenced *S. erinaceieuropaei* genome. The *S. erinaceieuropaei* genome was sequenced previously [12], with an estimated genome size of more than 1.2 Gb, but because the source material was from a biopsy the assembled sequence was highly fragmented. Hence, the *S. erinaceieuropaei* genome presented herein provides a more reliable estimate of the size and contents of this parasite genome. The new genome assembly was about two thirds of the size of the previous assembly but remains the largest genome among sequenced tapeworms. Compared to cyclophyllidean tapeworms, including *Echinococcus* and *Taenia* spp., for which high-quality genome references are available [16, 29, 30], genome information for pseudophyllidean tapeworms is limited [31]. The genomes presented in this study could, therefore, serve as a powerful resource for more comprehensive studies of tapeworm genomics and will facilitate the understanding of pseudophyllidean tapeworm biology and parasitism.

There have been three big knowledge gaps for the present cryptic tapeworm: 1) its phylogenetic relationship with *Spirometra* species, 2) its lifecycle including the definitive and intermediate hosts, and 3) genetic and physiological differences with non-proliferating *Spirometra* species that enable the worm to reproduce asexually in non-definitive hosts, such as humans and mice.

To determine phylogenetic relationships, we confirmed that the genetic sequence of *S. proliferum* is distinct from that of *S. erinaceieuropaei*, despite the close relationship between these species. Specifically, the *S. proliferum* genome is about 150-Mb smaller and contains 5000 fewer protein coding genes than in *S. erinaceieuropaei*. Both genomes, nonetheless, showed diploidy. These data suggest that *S. proliferum* is not an aberrant form of *Spirometra* worm by virus infection or by small mutations [6, 7] and not a hybrid origin of multiple *Spirometra* species. In agreement, no virus-like sequences were detected in *S. proliferum* DNA or RNA raw reads.

We were unable to identify definitive or intermediate hosts of *S. proliferum* in the current study. Recent horizontal transfers of genes or mobile elements can indicate phylogenetic relationships, because HGT events occur between closely associated organisms. Well-known examples include HGT from Wolbachia symbionts to their host insect [32, 33] and transfer of BovB retrotransposons between ruminants and snakes via parasitic ticks [34, 35]. We found that RTE/BovB repeats are abundant in the *S. proliferum* genome, but were likely acquired by an ancestral pseudophyllidea, as indicated by their abundance in *D. latum* and *S. solidus*. Moreover, our HGT screening analyses indicate several genes that were likely acquired from bacteria but these HGTs likely have occurred before specification of *S. proliferum* and *S. erinaceieuropaei*. The high-quality reference genomes presented herein, however, provide valuable resources for further attempts to identify vestigial *S. proliferum* sequences in other organisms or to perform analyses of protein–protein interactions between hosts and parasites.

Loss of genes that are involved in the development of multicellular organisms and nervous systems, including homeobox genes and genes for zinc-finger domain containing proteins, and relaxed selection of some developmental genes (ROBO, Slit, RHOGAP, etc.) suggests that *S. proliferum* has lost the ability to undergo proper development and complete the sexual lifecycle. Although the precise functions of homeobox genes in tapeworms remain elusive, proteins of homeobox families that are missing in *S. proliferum* (TALE/Pknox, ANTP/Hox1, ANTP/Msxlx and POU/Pou-like) appear to have important roles in the development of embryos and adult body plans. For example, Hox1 of the HOX gene family specifies regions of the body plan of embryos and the head–tail axis of animals [36]. Products of the Pknox gene family, also known as the PREP gene family, are implicated as cofactors of Hox proteins [37]. M*sxlx* homeobox gene was highly upregulated in the ovaries and was continually expressed in fertilized ova in the uterus in *Hymenolepis microstoma*. This gene was related to the female reproductive system in this tapeworm [38]. POU class genes are present in all animals and are extensively in nervous system development and the regulation of stem cell pluripotency in vertebrates [39]. Specific loss of Pou-like genes and relaxed selection of Pou3 suggest that *S. proliferum* has low dependency on POU genes.

We contend that the loss of sexual maturity of this parasite is related to its fatal pathogenicity in humans, because survival of the parasite is dependent on asexual reproductive traits of budding and branching, which lead to 100% lethality in infected humans. Accordingly, we identified genes that are upregulated in vigorously budding worms using transcriptome analyses and then selected genes that are putatively important for asexual proliferation, such as a variety of peptidase genes and oncogene-like genes. Among them, groups of secreted proteins with unknown functions were of great interest. They were expanded in the *S. proliferum* genome and showed more than 10-fold changes in expression levels. Recently, an *S. erinaceieuropaei* gene belonging to one of those groups (orthogroup OG0000083) was cloned and named plerocercoid-immunosuppressive factor (P-ISF) (Yoko Kondo, under review). P-ISF is a cysteine-rich glycoprotein abundant in plerocercoid excretory/secretory products and likely involved in immunomodulation of its hosts by suppressing osteoclastgenesis including the gene expression of TNF-α and IL-1β, and nitric oxide production in macrophages [40, 41]. Upregulation of P-ISF genes in *S. proliferum* proliferating worms is therefore reasonable and the expansion of the gene family in *S. proliferum* indicates the considerable contribution to the specific lifecycle. The other upregulated gene families of unknown function are also expanded in *S. proliferum* suggesting possible important roles in the hosts, therefore, future studies of these novel genes are required to fully understand the mechanism underlying the *S. proliferum* parasitism.

Fibronectin is an extracellular matrix (ECM) glycoprotein that controls the deposition of other ECM proteins, including collagens and latent TGF-beta binding protein [42]. During branching morphogenesis, accumulations of fibronectin fibrils promote cleft formation by suppressing cadherin localization, leading to loss of cell–cell adhesion [43]. The present observations of the *S. proliferum* lineage show specific expansions of three gene families containing fibronectin type III domains. *S. proliferum* also had fewer cadherin genes than *S. erinaceieuropaei* and three of them are subject to relaxed selection in *S. proliferum*. These results collectively suggest nonordinal ECM coordination in *S. proliferum*, allowing the formation of highly branching structures and enabling asexual proliferation in the host.

## Methods

### Biological materials

*S. proliferum* strain Venezuela was used for the genome analyses. The parasite was originally isolated from a Venezuelan patient in 1981 and has been maintained by serial passages using BALB/c mice via intraperitoneal injections of the plerocercoids in National Science Museum as described in Noya et al [44, 45]. *S. erinaceieuropaei* was isolated from a Japanese four-striped rat snake (*Elaphe quadrivirgata*) collected in Yamaguchi prefecture, Japan in 2014.

### DNA and RNA extraction and sequencing

*S. proliferum* worms were collected from the abdominal cavity of infected mice and washed thoroughly with 1x PBS. Plerocercoids of *S. erinaceieuropaei* were isolated from the subcutaneous tissues of the snake. Genomic DNA was extracted using Genomic-tip (Qiagen) following the manufacturer’s instructions.

Paired-end sequencing libraries (Additional Table 1) were prepared using the TruSeq DNA Sample Prep kit (Illumina) according to the manufacturer’s instructions. Multiple mate-paired libraries (3, 8, 12 and 16 kb) were also constructed using the Nextera Mate-Paired Library Construction kit (Illumina). Libraries were sequenced on the Illumina HiSeq 2500 sequencer using the Illumina TruSeq PE Cluster kit v3 and TruSeq SBS kit v3 (101, 150 or 250 cycles × 2) or the Illumina MiSeq sequencer with the v3 kit (301 cycles × 2) (Additional Table S1). The raw sequence data were analysed using the RTA 1.12.4.2 analysis pipeline and were used for genome assembly after removal of adapter, low quality, and duplicate reads.

RNA was extracted from individual worms using TRI reagent according to the manufacturer’s instructions. Total RNA samples were qualified using Bioanalyzer 2100 (Agilent Technology, Inc.). Only samples with an RNA integrity value (RIN) greater than 8.0 were used for library construction. One hundred ng of total RNA was used to construct an Illumina sequencing library using the TruSeq RNA-seq Sample Prep kit according to the manufacturer’s recommended protocols (Illumina, San Diego, USA). The libraries were sequenced for 101 or 151 bp paired-ends on an Illumina HiSeq2500 sequencer using the standard protocol (Illumina).

### K-mer Analysis

A k-mer count analysis was performed using K-mer Counter (KMC) [46], on the paired-end Illumina data. Only the first read was used to avoid counting overlapping k-mers. Genome size and ploidy estimations were performed using Genomescope [47] and Smudgeplot, respectively [48].

### Genome assembly

Illumina reads from multiple paired-end and mate-pair libraries (Additional Table 1) were assembled using the Platanus assembler [49] with the default parameter. Haplomerger2 [50] was then used to remove remaining haplotypic sequences in the assembly and contigs were further scaffolded using Illumina mate-pair reads using SSPACE [51]. CEGMA v2 [52] and BUSCO [53] were used to assess the completeness of the assemblies.

Mitochondrial genomes (mitogenomes) were reconstructed from Illumina reads with MITObim version 1.6 [54]. Mitochondrial fragments in the nuclear genome assembly were identified by BLASTX using *S. mansonai* mitochondrial genes as queries and those fragments were extended by iterative mappings of Illumina short reads using MITObim. Assembled mitogenomes were annotated for protein-coding, tRNA and rRNA genes using the MITOS web server [55]. Assemblies and annotations were manually curated using the Artemis genome annotation tool [56] with based on evidence supports from sequence similarity to other published mitogenomes.

### Repeat analysis

Repeats within the genome assemblies were identified using RepeatModeler (v1.0.4, http://www.repeatmasker.org/RepeatModeler.html) and RepeatMasker (v.3.2.8, http://www.repeatmasker.org) to calculate the distribution of each repeat and its abundance in the genome.

### Gene prediction and functional annotation

To predict protein-coding genes, Augustus (v. 3.0.1) [57] was trained for *S. proliferum* and *S. erinaceieuropaei*, individually, based on a training set of 500 non-overlapping, manually curated genes. To obtain high-confidence curated genes, a selection of gene models from gene predictions based on Augustus *S. mansonai* parameters, were manually curated in Artemis using aligned RNA-seq data and BLAST matches against the NCBI database. RNA-seq reads were mapped to the genomes using Hisat2 (parameters: --rna-strandness RF --min-intronlen 20 –max-intronlen 10000) [58]. Based on the Hisat2 alignments, the bam2hints program (part of the Augustus package) was used to create the intron hints, with minimum length set to 20 bp. Augustus were run with trained parameters using all the hints for that species as input. Introns starting with ‘AT’ and ending with ‘AC’ were allowed (--allow_hinted_splicesites=atac). A weight of 10^5^ was given to intron and exonpart hints from RNA-seq. If Augustus predicted multiple, alternatively spliced transcripts for a gene, we only kept the transcript corresponding to the longest predicted protein for further analyses.

Functional annotations were performed on the gene models based on multiple pieces of evidence including BLASTP search against NCBI nr database and the latest version Pfam search (ver. 30.0) with HMMER3 [59]. Gene ontology (GO) terms were assigned to genes using Blast2Go (v2) [60] with BLAST search against NCBI nr database and the InterProScan results.

### Species tree reconstruction

Amino acid sequences in each single-copy gene family were aligned using MAFFT version v7.22152 [61], poorly aligned regions were trimmed using GBlocks v0.91b53 [62], and then the trimmed alignments were concatenated. A maximum-likelihood phylogenetic tree was produced based on the concatenated alignment using RAxML v8.2.754 [63] with 500 bootstrap replicates. The best-fitting substitution model for each protein alignment was identified using the RAxML option (-m PROTGAMMAAUTO). Mitochondrial genome phylogeny was also constructed by the same method using 12 protein coding genes on mitogenomes.

### Gene family analysis

To estimate branch or lineage specific gain and loss of orthologous gene families, OrthoFinder [64] and CAFÉ (v3) [65] under parameters “-p 0.01, -r 1000” were used.

### Screening for horizontally transferred genes

To screen potential horizontal gene transfers (HGTs) into the *S. proliferum* and *S. erinaceieuropaei* lineages, we used DarkHorse v2, which detects phylogenetically atypical proteins based on phylogenetic relatedness of blastp hits against a taxonomically diverse reference database using a taxonomically-weighted distance algorithm [66]. Options (-n 1 -b 0.5 -f 0.1) were used in DarkHorse HGT screening.

### Positive Selection Scans (dNdS)

To analyse selection pressures in *S. proliferum* genes, the ETE3 Python package [67] for CODEML [68] was employed to calculate the non-synonymous (dN) and synonymous (dS) substitutions rates, and the ratio (dN/dS or ω). Nucleotide sequences of single copy orthologue genes from 12 cestode species (*S. proliferum*, *S. erinaceieuropaei*, *Diphyllobothrium latum, Schistocephalus solidus, Hymenolepis diminuta, Hymenolepis nana, Hydatigera taeniaeformis, Taenia solium, Taenia asiatica, Echinococcus multilocularis, Echinococcus granulosus, Mesocestoides corti*) were aligned based on amino acid alignment using Pal2aln v14 [69] with the parameters (- nomismatch and –nogap). dN/dS were estimated using branch-site models with *S. proliferum* as the foreground and other branches in the tree as the background. The non-null model (bsA) were compared with the null model (bsA1) for each tree using a likelihood ratio test (LRT), where log-likelihood ratios were compared to a chi-square distribution with 1 degree of freedom. False discovery rate (FDR) correction were performed over all the P-values and genes showing FDR <0.05 were manually curated before obtaining final dN/dS values.

Test for relaxed selection was performed using the RELAX tool [70] with aforementioned single copy orthologue gene sets. The relaxation parameter k was calculated for each blanch and tested by LRT with *S. proliferum* as foreground and the others as background.

### RNAseq analysis

For gene expression analyses, *S. proliferum* plerocercoid worms were grouped into two types based on the morphology and proliferation activity; worms vigorously branching to form structure like “Medusa head” and worms under static form to form like “Wasabi root” (Figure 4a). Worms were collected from infected mice on ~50 weeks post inoculation. RNA was extracted from the individual worms and sequenced as described above. RNAseq reads were mapped to the *S. proliferum* reference genomes (v2.2) using Hisat2 [58] (parameters: --rna-strandness RF --min-intronlen 20 –max-intronlen 10000). Mapped read count of each gene was calculated using HTSeq [71] with options (-s no, -a 10, -m union) and differential expression analyses were performed using EdgeR v3.2.4 [72]. A transcript was identified as differentially expressed in a pairwise comparison if the following criteria were met: false discovery rate (FDR) ≤ 0.001 and fold change ≥ 2.0. FPKM values were calculated using Cufflinks packages v2.2.1 [73] and used to generate for multidimensional scaling (MDS) plots and gene expression heatmaps.

## Declarations

### Ethics approval and consent to participate

All animal experiments in this study were performed under the applicable laws and guidelines for the care and use of laboratory animals, as specified in the Fundamental Guidelines for Proper Conduct of Animal Experiment and Related Activities in Academic Research Institutions under the jurisdiction of the Ministry of Education, Culture, Sports, Science and Technology, Japan, 2006.

### Consent for publication

Not applicable.

### Availability of data and materials

All sequence data from the genome projects have been deposited at DDBJ/ENA/GenBank under BioProject accession PRJEB35374 and PRJEB35375. All relevant data are available from the authors.

### Authors’ Contributions

T.Ki., T.Ku. and H.M. conceived the study. T.Ki. contributed to study design. V.L.H., H.M. and T.Ki. wrote the manuscript with inputs from others. BAN, ON, SK prepared biological samples. Ki and T.Ku. conducted experiments. V.L.H., M.D., Y.M., A.T. and T.Ki. completed genome assembly and analysed genome data.

## Acknowledgements

Genome data analyses were partly performed using the DDBJ supercomputer system. We thank Ryusei Tanaka, Asuka Kounosu, Akemi Yoshida for assistance and comments.

## Author Information

Correspondence and requests for materials should be addressed to T.K.

