## Supplementary figures for "Genome analysis of the fatal tapeworm *Sparganum proliferum* unravels the cryptic lifecycle and mechanisms underlying the aberrant larval proliferation"

**This PDF file includes:**
Additional Fig S1 to S5

**Other Supplementary Materials for this manuscript includes the following:**
Additional Table S1 – S14 (an excel file)

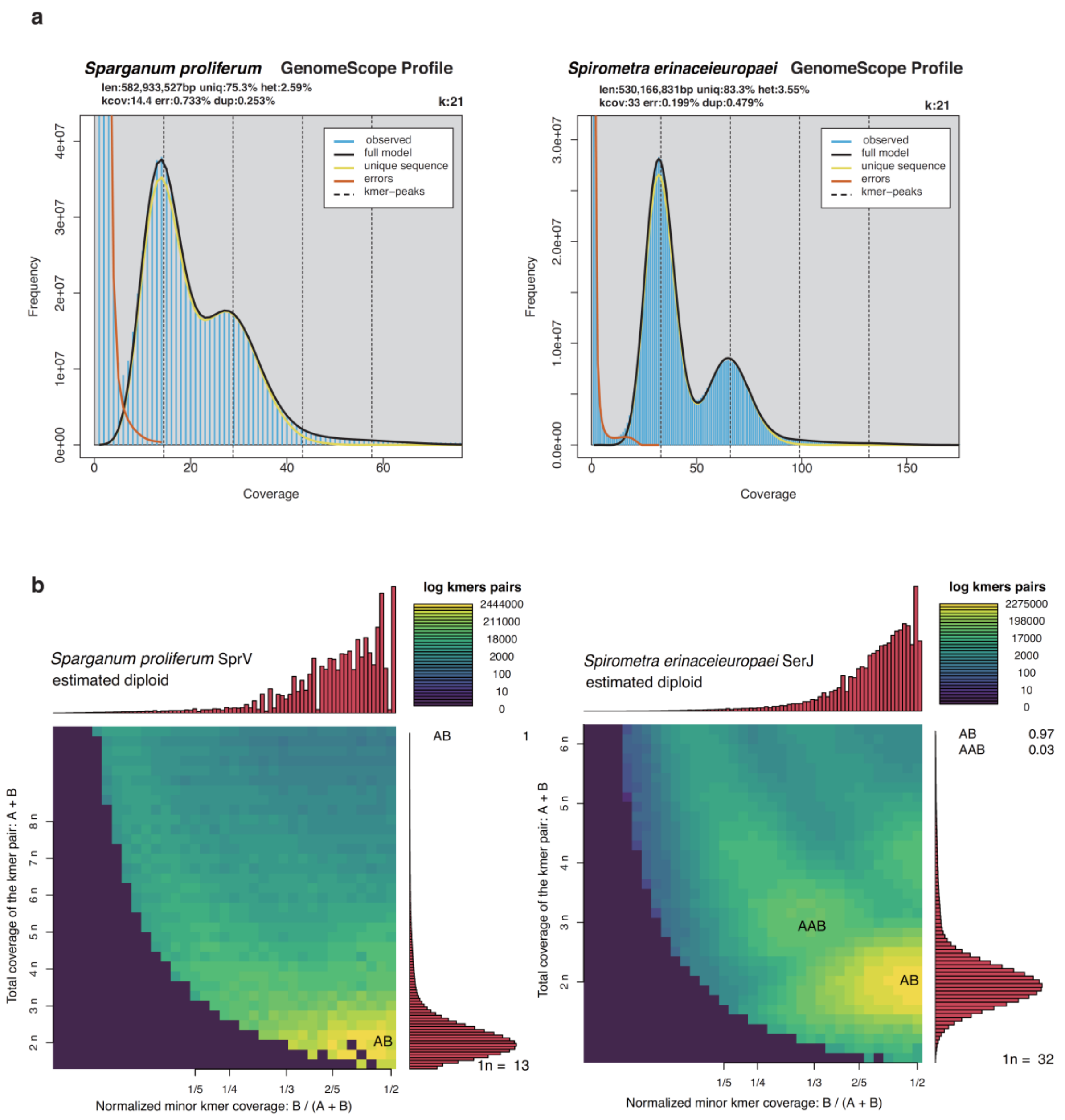


Additional Fig S1. (a) Histograms of the 23-mer depth distribution for *S. proliferum* and *S. erinaceieuropaei* were plotted by GenomeScope to estimate genome sizes, repeat contents, and heterozygosity levels. (b) Ploidy was estimated using Smudgeplots for *S. proliferum* and *S. erinaceieuropaei*.

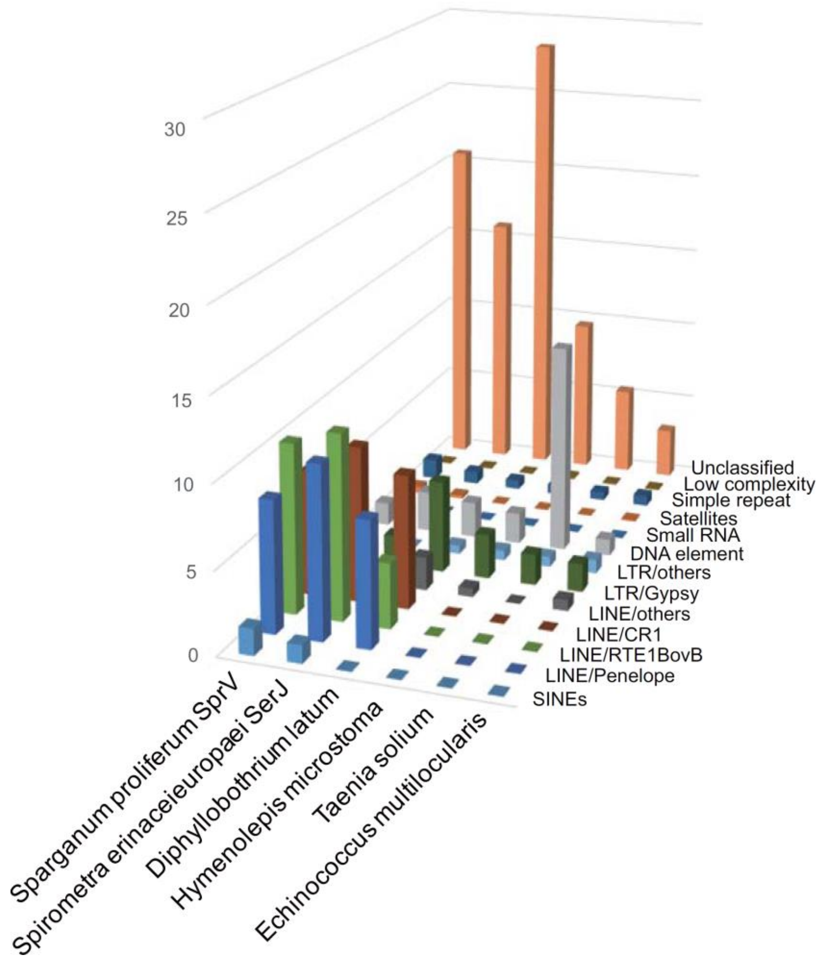


Additional Fig S2. Comparison of genome repeat contents among cestode species

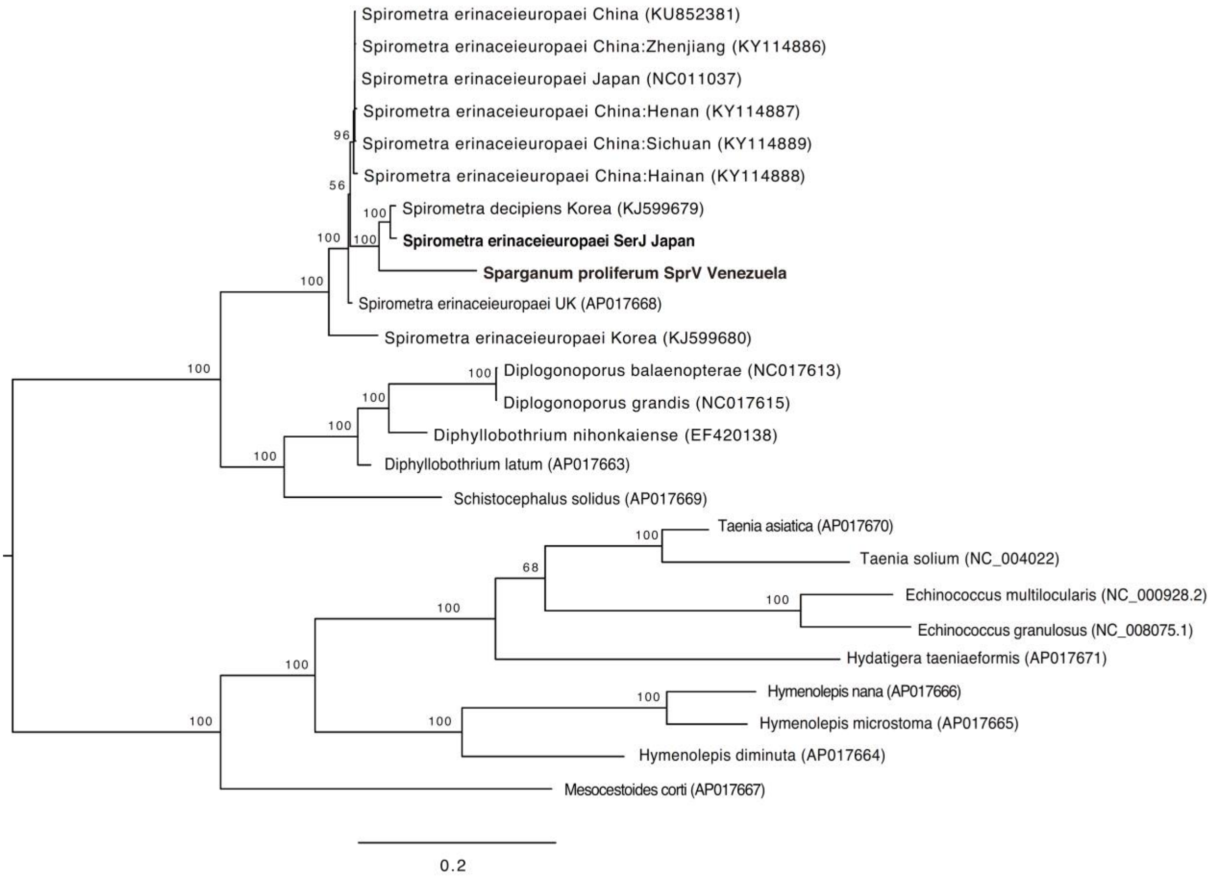


Additional Fig S3. Maximum-likelihood phylogenetic tree based on the mitochondrial genomes (12 protein-coding genes) of cestodes; amino acid sequences were aligned and phylogenetic analyses were performed with RAxML using the best-fitting empirical model of amino acid substitution with 1,000 bootstrap resampling replicates and the percentage support shown on the nodes. The scale bar shows the number of amino acid substitutions per site.

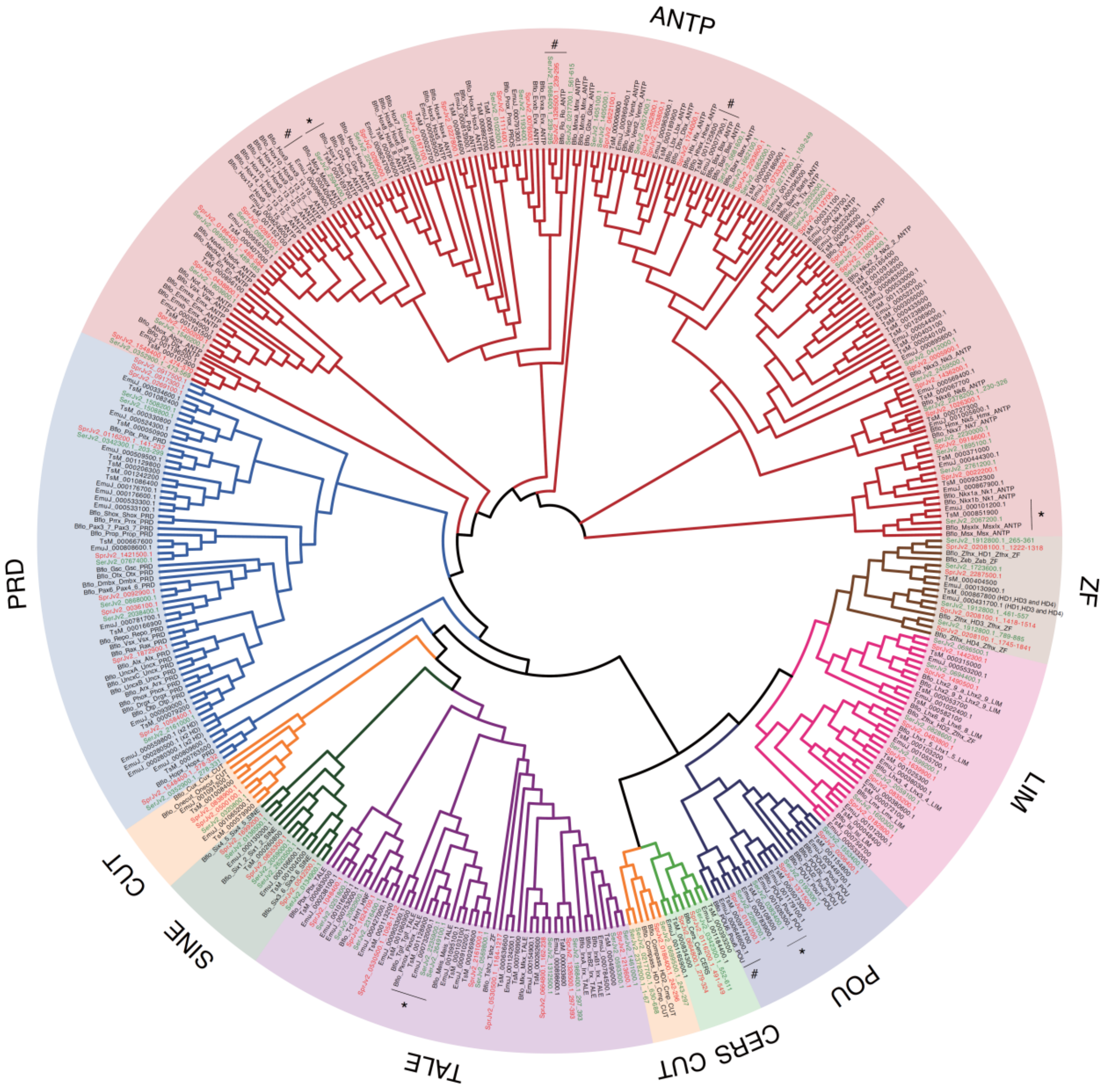


Additional Fig S4. Cladogram showing the diversity of homeobox genes in *S. proliferum* and *S. erinaceieuropaei* with the tapeworm species *T. solium* and *E. multilocularis* and the bilaterian *Branchiostoma floridae*.

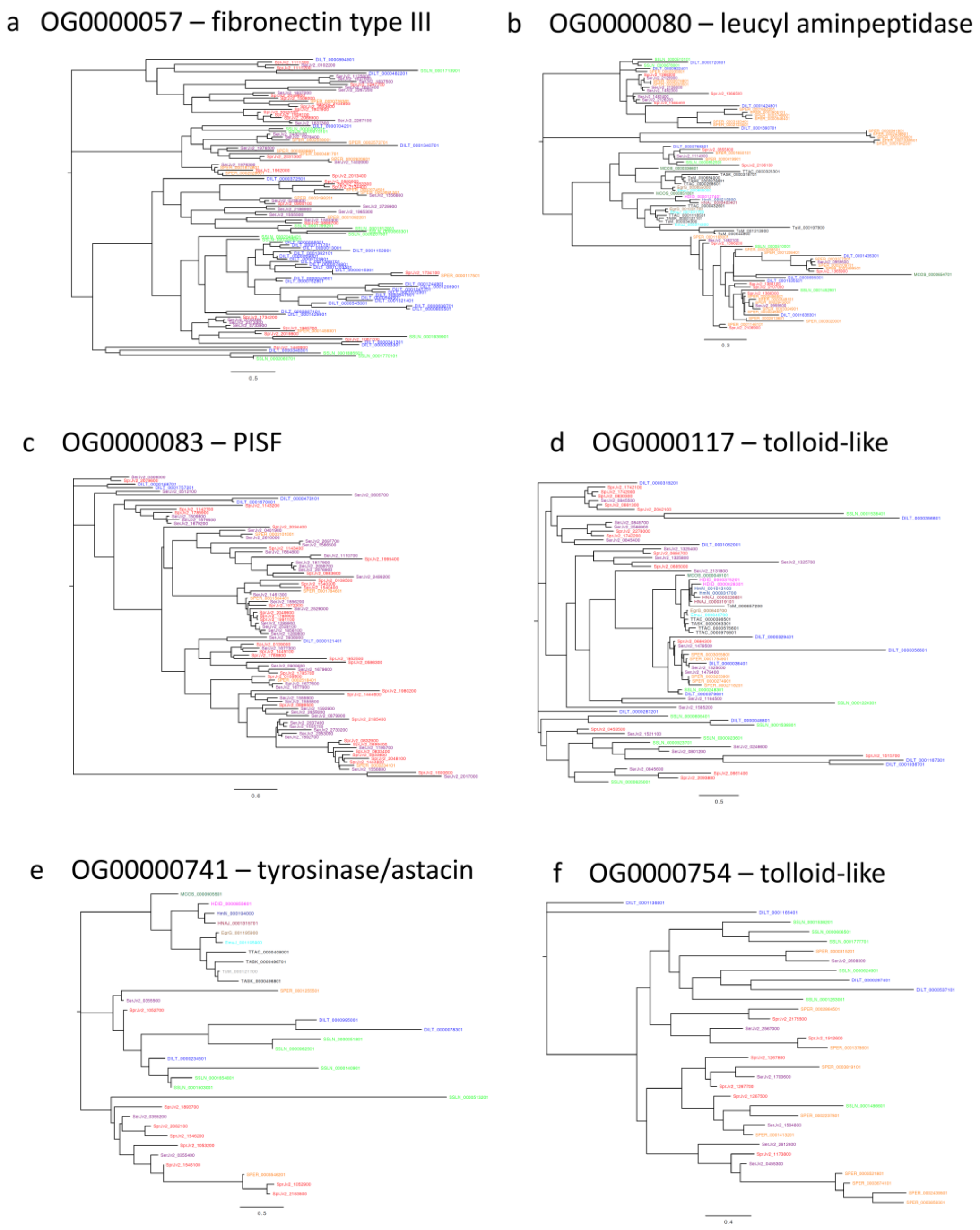


Additional Fig S5. Phylogenetic trees for six orthofamilies that are differentially expressed between the medusa-head and wasabi-root forms of *S. proliferum*. Orthofamilies have been identified by Orthofinder analysis and include 12 species (*S. proliferum* = red, *S. erinaceieuropaei* =purple, *Diphyllobothrium latum* =blue, *Schistocephalus solidus* = green, *Hymenolepis diminuta* = magenta, *Hymenolepis nana* = dark red, *Hydatigera taeniaeformis* = black, *Taenia solium* = grey, *Echinococcus multilocularis* = cyan, *Echinococcus granulosus* = brown, *Mesocestoides corti* = dark green). Phylogenies indicate that some of these orthofamilies are conserved across flatworms (e.g. a), while others are specific to the Pseudophillidea clade of flatworms (e, g and f). The scale bar represents amino acid substitutions.
